# KCC2 as a novel biomarker and therapeutic target for motoneuron degenerative disease

**DOI:** 10.1101/2023.08.24.554410

**Authors:** C. Sahara Khademullah, Julien Bourbonnais, Mathilde M. Chaineau, María José Castellanos-Montiel, Iason Keramidis, Alexandra Legault, Marie-Ève Paquet, Agessandro Abrahao, Lorne Zinman, Janice Robertson, Thomas M. Durcan, Melanie A. Woodin, Antoine G. Godin, Yves De Koninck

## Abstract

Hyperexcitability in cells throughout the corticospinal tract is a presymptomatic feature of amyotrophic lateral sclerosis (ALS) associated with lethal motor degeneration ^1–6^. Disinhibition is a possible cause of this hyperexcitability, potentially implicating the central nervous system-specific potassium-chloride cotransporter, KCC2, a core regulator of the strength of GABAergic neurotransmission linked to several neurological disorders ^7–11^. Here, we show that KCC2 is downregulated in the membrane of motor cortex neurons from post-mortem SOD1-, C9orf72- and sporadic ALS is patients. Increased protein levels of KCC2 were found in plasma and cerebral spinal fluid of ALS patients and mice harbouring the SOD1*G93A mutation. Longitudinal analysis of disease progression in both SOD1*G93A and Prp-TDP43*A315T mice revealed a decrease of KCC2 membrane levels in cortical and spinal motor neurons which were already present at the presymptomatic phase. Using KCC2-enhancing compounds, CLP290 and prochlorperazine (PCPZ) restored KCC2 membrane expression and function, delayed motor deficit onset, and extended lifespan up to two months in mutant mice. Human-derived neurons differentiated from iPSC harbouring the SOD1*G93A mutation displayed KCC2 deficits which PCPZ treatment rescued. Acute administration of KCC2 enhancers restored chloride transport in presymptomatic and symptomatic mice and reversed motor neuron hyperexcitability in awake behaving mutant mice. These findings identify KCC2 as both an early biomarker and a disease-modifying therapeutic target for ALS.

## Main

Amyotrophic lateral sclerosis (ALS) is a rapidly progressive, fatal neurodegenerative disorder that preferentially affects both the upper and lower motor neurons (UMN/LMN) in the central nervous system (CNS) ^12,13^. It is considered the most common form of motor neuron disease (MND), accounting for 80 – 90% of cases ^14^. The incidence of ALS across Europe and North America is approximately 2.5 - 3 cases per 100,000 people per year ^15^. Following the initial onset of symptoms, approximately 50% of cases are fatal within the first 1.5 years and in about 20% of cases, the disease persists for approximately 5 to 10 years, often resulting in respiratory failure ^16,17^. The etiology of most cases of ALS is unknown, with more than 90% of ALS cases classified as sporadic ALS (sALS), and the remaining falling into the category of familial ALS (fALS) ^18^. Given the lack of diagnostic biomarkers and the complete absence of curative treatments, identifying treatable targets that can also serve as diagnostic biomarkers that are present across all ALS cases is urgently required.

A balance between synaptic excitation and inhibition is essential for normal brain function. When this delicate balance is disrupted, it can lead to neuronal hyperexcitability ^7–10^. Electrophysiological recordings from ALS patients and animal models of the disease have revealed an early disruption to this balance ^1,3^. In fact, cortical hyperexcitability is one of the earliest features of the disease and it has been suggested that it can potentially propagate through the corticomotor system causing deficits in neuronal transmission and eventual degeneration of lower motor neurons ^19–22^. In line with this, clinical transcranial magnetic stimulation (TMS) studies from patients carrying either SOD1 or C9orf72 mutations have provided evidence that deficits in cortical inhibition precede the onset of cortical hyperexcitability ^23^. Early decreases in cortical inhibition occurring ahead of increases in excitation have also been identified by us and others in mouse models of the disease to further support this hypothesis ^1,6,19,24,25^. While the underlying substrates of this altered state of inhibition can be vast, recent studies have identified a particularly interesting new target to restore activity levels: the central nervous system (CNS)-specific potassium (K^+^)-chloride (Cl^-^) cotransporter, KCC2. Indeed, decreased KCC2 gene expression and neuronal membrane expression in cultured spinal cord motor neurons has been reported in the SOD1*G93A mouse model of ALS^26^. Taken together, these findings prompted us to ask if deficits in KCC2 expression are present in ALS patients and animal models carrying ALS-linked mutations. We also asked whether plasma and CSF levels of KCC2 could serve as proxies of altered KCC2 expression in the CNS and thus as a potential biomarker of the disease.

## Results

### KCC2 expression is reduced in the motor cortex of ALS patients and enhanced in plasma and CSF

Clinical transcranial magnetic stimulation studies of the primary motor cortex (M1) from presymptomatic and symptomatic patients carrying known ALS mutations have revealed a loss of short-cortical inhibition ^2,6,22–24^. Since KCC2 is a key regulator of inhibition in the CNS ^27^, we first hypothesized that its expression would be altered in M1 tissue from post-mortem ALS patients. Using immunofluorescence imaging, we assessed KCC2 expression at the plasma membrane of neurons in the layer 5 of the motor cortex (L5-M1) ^28^ taken from three groups of post-mortem ALS patients with either sporadic ALS, SOD1 or C9orf72 mutations. Our results revealed a significant decrease in KCC2 expression in neuronal membranes across all three patient groups (**Fig. 1a - d**), indicating that deficits in KCC2 expression may be a common feature across sALS and fALS cases. This decrease in KCC2 membrane expression prompted us to investigate whether a corollary of this was detectable in the plasma or cerebral spinal fluid (CSF) from recently diagnosed sporadic ALS patients. Results using a customized sandwich Elisa kit specific for the C-terminus of KCC2 indicated that protein levels were significantly increased in plasma serum and CSF from ALS patients (**Fig. 1e**). Given that our immunofluorescent data revealed a loss of internalized and membrane bound KCC2 expression (**Fig. 1c**), membrane shedding of the protein may be culpable in the observed increase in the biological fluids. Together, these finding reveal that changes in KCC2 -membrane expression in the M1 and -protein levels in the blood and CSF are common features observed in several forms of ALS. Moreover, this data provides the first evidence that KCC2 protein levels may potentially serve as a biomarker in ALS.

**FIGURE 1:**
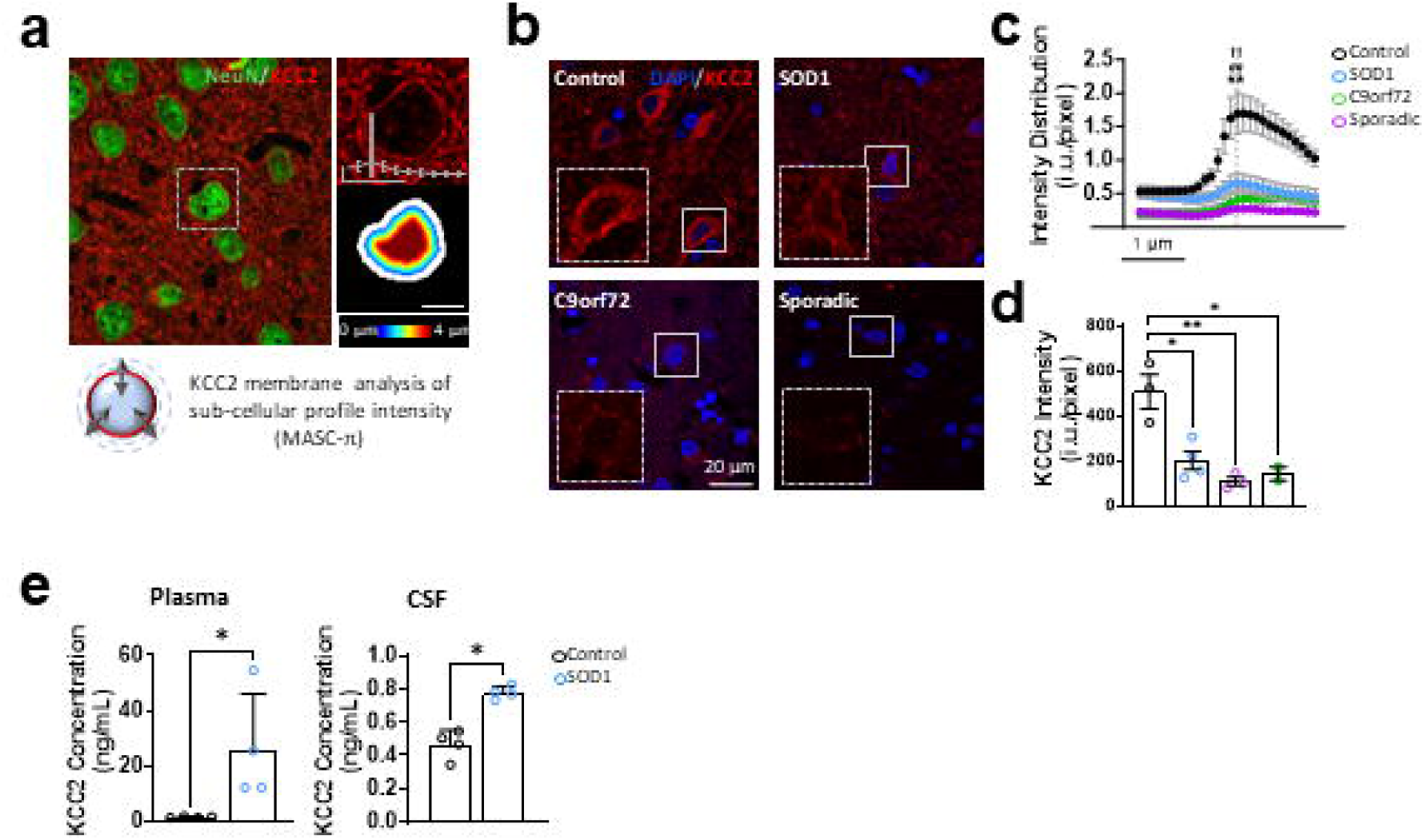
KCC2 expression is reduced in the motor cortex of ALS patients and enhanced in plasma and CSF. **(a)** Example confocal image with color-coded distance map illustrating profile intensities (MASC-⫪). **(b)** Representative confocal images from the L5-M1 of healthy individuals *(N* = 3) and ALS patients with known SOD1 *(N = 4),* C9orf72 *(N* = 2) or Sporadic *(N = 3)* mutations. **(c)** Quantification of KCC2 intensity as a the shift in KCC2 intensity with the distance to the membrane profile Inset shows membrane analysis of sub-cellular function of distance to the membrane. Dotted grey line denotes the membrane. Asterisks denotes significance for SOD1 Hashtags denote significance for sporadic. Daggers denote significance for C9orf72. **(d)** Quantification of KCC2 intensity at the membrane. Unpaired *t*-test. **(e)** Quantification of KCC2 in plasma and CSF extracted from newly-diagnosed sporadic ALS patients with a SOD1 mutation. *N = 4.* Student *t*-test with Mann Whitney post-hoc. Data are presented as mean ± SEM. * P ≤ 0.05. ** P ≤ 0.01

### Reduced expression of KCC2 is specific to the motor cortex and lumbar ventral horn of the asymptomatic SOD1 and TDP43 mutant mice, while enhanced in CSF

To both investigate the temporal relationship between the different stages of the disease pathology and KCC2 expression and confirm our results acquired from ALS patients, we asked whether a deficit in KCC2 protein expression could be observed in mice harbouring the SOD1*G93A mutation. To date, SOD1-ALS-causing mutations have been widely studied and account for some 20% of familial ALS and another 5% of sporadic ALS cases ^29^. While controversial for its high copy numbers of mutant SOD and rapid degeneration, the familial SOD1*G93A mutant mouse model remains the gold standard for motor deficit assessment associated with ALS ^30,31^. To investigate the temporal relationship between the different stages of the disease pathology and KCC2 expression, we asked whether a deficit in KCC2 protein expression could be observed in SOD1*G93A mutant mice. We probed tissue containing the L5-M1 for KCC2 during the asymptomatic (P30), presymptomatic (P60) and symptomatic stages (P100 - 115) of ALS and found that membrane KCC2 expression was already decreased in asymptomatic SOD1 mutant mice, which also coincides with the juvenile stage of development (**Fig. 2a, b & d**). Since ALS is a disease that predominantly affects not only cortical motor neurons, but spinal motor neurons as well ^32^, we probed the L5 segment of the spinal cord (L5-SC) and found a similar decrease in KCC2 expression in the ventral horn (VH) (**Fig. 2a, b & d**). TAR DNA-binding protein 43 (TDP43) pathology can be found in as many as 97% of sporadic and familial cases of ALS, making this model highly clinically relevant ^33–37^, thus we asked whether the KCC2 deficits were also present in TDP43*A315T mutant mouse. Our results indicated that KCC2 expression was significantly decreased in both the L5-M1 and L5-VH of TDP43 mutant mice as early as the presymptomatic stage continuing through to the symptomatic stage (**Fig. 2c & d**). The downregulation of KCC2 at the neuronal membrane in SOD1 and TDP43 mutant mice did not coincide with any changes in the overall NeuN^+^ cell count nor MAP2 expression in SOD1 mice, indicating that a reduction in KCC2 expression is specific and not the result of cell loss (**Fig. 2e & f**). Membrane KCC2 expression was also unchanged in the Nucleus Accumbens of SOD1 mice when compared to their non-transgenic littermates, indicating that the loss observed was specific to the cortical and spinal motor areas (**Fig. 2g**). Finally, our Elisa assay revealed that there was a significant increase in KCC2 levels already in CSF extracts from presymptomatic SOD1 mutant mice when compared to their non-transgenic littermates (**Fig. 2h**). While KCC2 is specifically expressed in CNS neurons, detection of its expression levels in humeral compartments outside the CNS may serve as an early biomarker of the disease ^38^.

**FIGURE 2.**
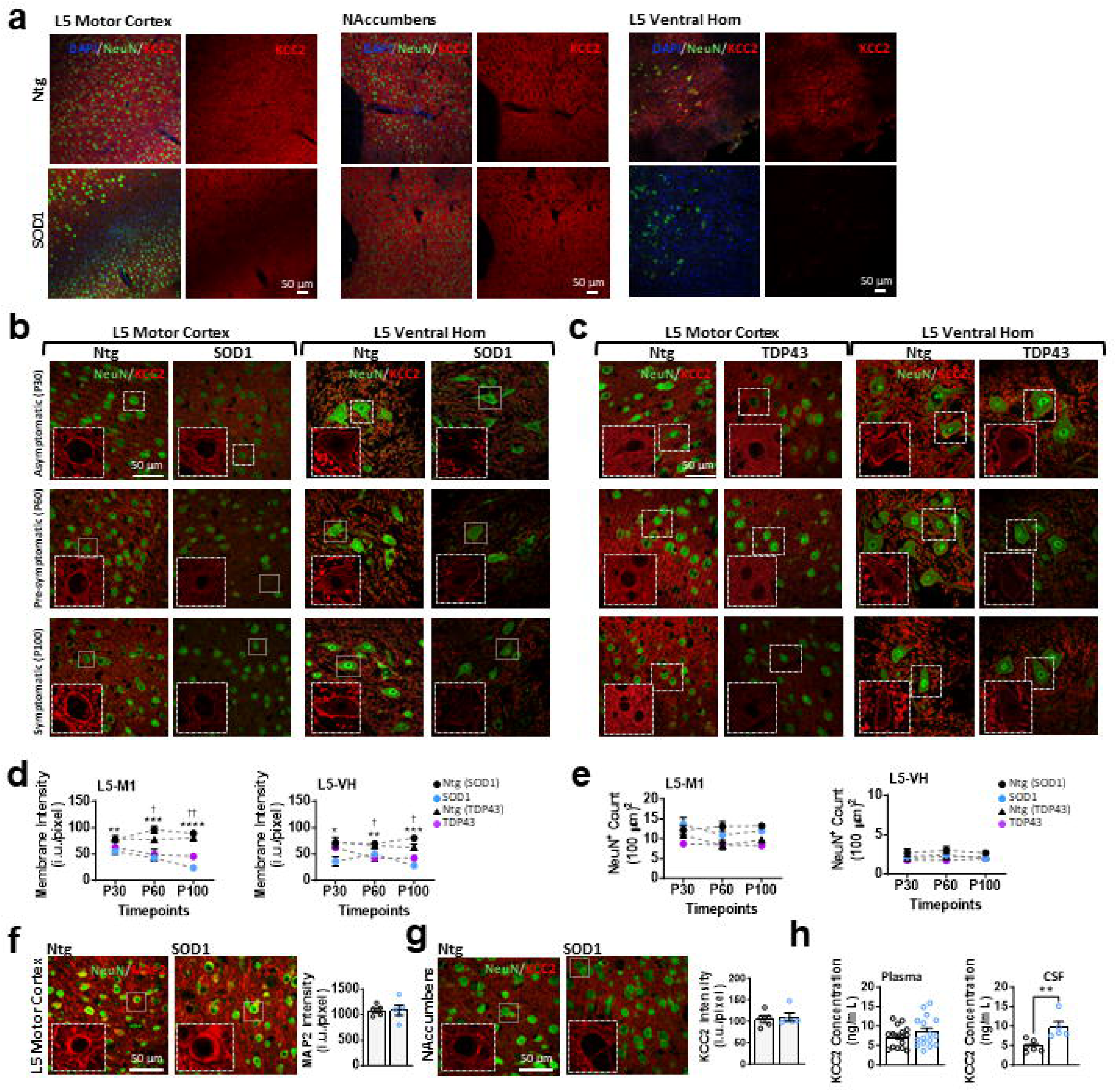
Reduced expression of KCC2 is specific to the motor cortex and lumbar ventral horn of the asymptomatic SOD1 and TDP-43 mutant mice, while enhanced in CSF. **(a)** Representative confocal images taken at 20X of DAPI, NeuN and KCC2 from L5-M1, nucleus accumbens. and the L5-VH from Ntg and SOD1 mice at P115. **(b)** Representative confocal images of NeuN and KCC2 from L5-M1 and the L5-VH from SOD1 mice at P30. P60, and P100. **(c)** Similar to (b) but taken from TDP43 mice. **(d)** Quantification of mean KCC2 for SOD1 and TDP43 mice. **(e)** Similar to (d) but quantification for NeuN^+^ cells **(f)** Representative confocal images and quantification of KCC2 intensity at the membrane in slices taken from the nucleus accumbens from NTg and SOD1 mice at P100. **(g)** Similar to (f) but for MAP2 intensity in slices taken from the L5-MC. *(N = 6 mice),* **(h)** Quantification of KCC2 protein levels in plasma *(N = 16)* and CSF *(n = 6, 2-3 samples pooled together from 16 mice)* from SOD1 mice. Mann-Whitney test Data are presented as mean ± SEM. * P ≤ 0.05. ** P 0.01. P ≤ 0.001, **** P ≤ 0.0001.

### Long-term administration of CLP290 and PCPZ restores KCC2 expression, preserves motor neuron health and delays the onset of motor deficits in SOD1 mice

We recently identified the arylmethylidine family as a group of compounds that increase KCC2 activity with virtually no side effects. Within this group, the CLP257 prodrug, CLP290, was shown to enhance KCC2 activity and lower intracellular Cl^-^ levels in several disease models ^8,11,39,40^. Moreover, while the loss of KCC2 has been implicated in motor deficits, its upregulation at the membrane *via* these compounds has proven to be essential in the recovery of motor function after spinal cord injury ^41–43^. Since we found that KCC2 membrane expression was significantly reduced in both the M1 and spinal cord, we sought to restore its expression in these motor areas by long-term administration of CLP290, which we also anticipated would result in improved motor function in the well-characterized SOD1*G93A mouse. To determine whether this family of compounds could protect against the hallmark motor symptoms associated with the disease and in turn, prolong survival, we treated presymptomatic SOD1 mutant mice with CLP290 (100 mg/kg/day, p.o.). With no conclusive diagnostic biomarkers for ALS, it is common for patients to be symptomatic by the time they visit a clinic and receive a diagnosis. In fact, patients typically experience delays of 10 to 16 months between symptom onset and diagnosis ^44–46^. Therefore, it is imperative that therapeutic strategies be developed and validated as close to the symptomatic stage of the disease as possible to recapitulate clinical patient presentation. However, since our data revealed a significant symptomatic loss of KCC2 at the plasma membrane of more than ∼ 75%, we began therapeutic administration during the presymptomatic phase of the disease where expression was reduced by ∼ 45%, leaving substantial room for a rescue in expression. We found that long-term administration of CLP290 starting at the presymptomatic stage of the disease, significantly delayed the onset of motor deficits (measured via grip strength and rotarod performance) in SOD1 mice compared to vehicle-treated SOD1 mice (**Fig. 3a**). Moreover, CLP290-treated SOD1 mice significantly outlived their vehicle-treated counterparts by approximately 2.5 weeks (**Fig. 3c**). Recent reports have identified that the traditional antipsychotic phenothiazine derivatives, including the FDA-approved prochlorperazine (PCPZ), at sub-clinical doses, enhance KCC2 activity ^42,47^. Thus, in a separate cohort of SOD1 mice, we administered low-dose PCPZ (1 mg/kg/day i.p.) long-term, once again beginning at the presymptomatic stage, and found that it delayed motor deficits and extended lifespan in SOD1 mice by more than two months (**Fig. 3b & d**). Moreover, PCPZ treatment was able to prevent further weight loss, a well-known sustained consequence of ALS. (**Fig. 3b**).

**FIGURE 3.**
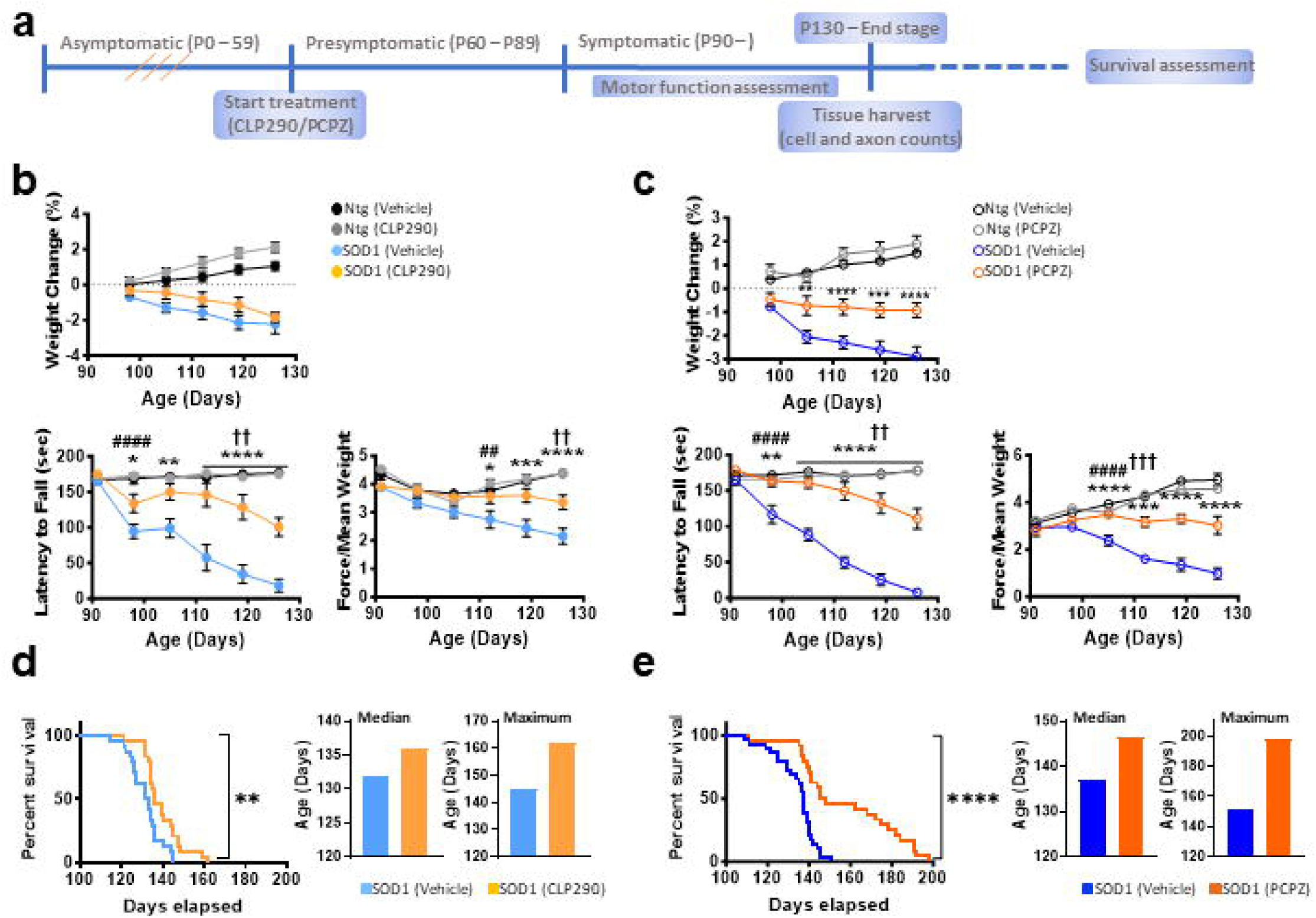
Long-term administration of CLP290 and PCPZ delays the onset of motor deficits and prolong survival in SOD1 mutant mice. **(a)** Schematic illustrating experimental timeline. **(b)** Weight and behavioural quantification of grip strength and rotorod tasks in mutant mice treated with CLP290 (100 mg/kg) *(N* = 24 for each group. 12 males and 12 females). Asterisks shown for SOD1 Vehicle and SOD1 CLP290 groups. Hashtags denote symptom onset between SOD1 and Ntg Vehicle groups. Daggers denote symptom onset between SOD1 CLP290 and Ntg Vehicle groups. **(c)** Same as (b) but with PCPZ (1 mg/kg) treatment. **(d)** Kaplan-Meier curve showing percent survival of SOD1 Vehicle and SOD1 CLP290 groups, with median and maximum survival graphs. **(e)** Similar to (d), but for SOD1 Vehicle and SOD1 PCPZ groups. (A/ = 24 for each group. 12 males and 12 females). Asterisks shown for SOD1 Vehicle and SOD1 treatment groups. Two-way ANOVA with Tukey’s multiple comparison post hoc. Data are presented as mean ± SEM. * P ≤ 0.05, “Pi 0.01, *** P ≤ 0.001, **** P ≤ 0.0001.

To determine whether treatment with the KCC2 enhancers were sufficient to restore KCC2 membrane expression and prevent neuronal loss, we quantified the number of motor neurons in the L5-M1 (**Fig. 4a**), L5-SC (**Fig. 4b),** as well as the number of L5 motor axons and measured KCC2 expression (**Fig. 4c**). We found that, following treatment, KCC2 expression was in fact restored at the neuronal membrane in both the L5-M1 (**Fig. 4a, d & f**) and the VH of the L5-SC (**Fig. 4b, e & g**). Not only was there a significant preservation of cortical (CTIP2^+^, **Fig. 4d**) and spinal motor neurons (NeuN^+^ cell >250 µm^2^, **Fig. 4e**), but there was also a preservation in the total number of cells compared to vehicle-treated SOD1 mice (total NeuN^+^, **Fig. 4d & e**). Moreover, PCPZ and CLP290 were able to significantly reduce the number of motor axons lost when compared to the vehicle treated SOD1 mutant mouse (**Fig. 4c, e, & g** and **Supplemental Fig. 1**).

**FIGURE 4.**
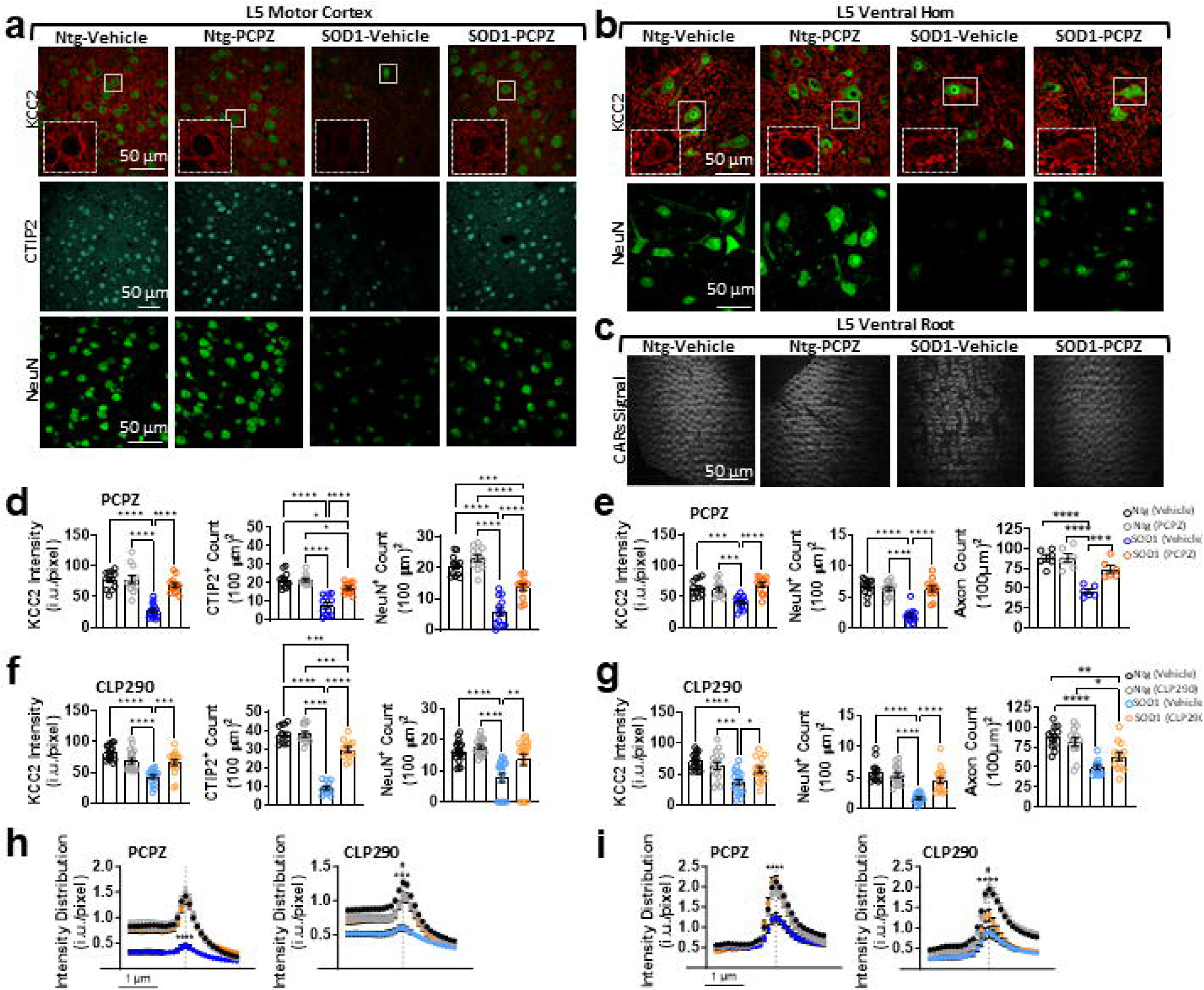
Long-term administration of CLP290 and PCPZ prevents KCC2 downregulation and delays neurodegeneration in S0D1 mutant mice. **(a)** Representative confocal images showing KCC2 membrane labelling. CTIP2^+^. and NeuN^+^ neurons in the L5-M1 at Week 18. **(b)** Similar to (a) but showing KCC2 membrane labelling and NeuN^+^ neurons in the L5-VH . **(c)** Images used to count the number of motor axons from the L5-VR taken with CARs **(d)** (left - right) Quantification of KCC2 intensity at the membrane, CTIP2^+^ and NeuN^+^ cell counts from the L5-M1 treated with PCPZ. **(e)** Similar to (d) but for KCC2 intensity at the membrane. NeuN^+^ cell counts from the L5-VH and motor axon counts from the L5-VR *(N = 12-14 forKCC2, CTIP2^+^ and NeuN^+^, N = 6 for VR count)*. **(f & g)** Similar to (d & e) but with CLP290 administration. *N = 12-18 for KCC2, CTIP2*^+^ *and NeuN^+^, N = 12 for VR count.* Also see Supplemental Fig 1. **(h)** Quantification of KCC2 intensity as a function of distance to the membrane in the presence of PCPZ (left) and CLP290 (right) in the L5-M1. **(i)** Similar to (h) but in the L5-VH. Dotted grey line denotes membrane. Asterisks denotes significance between vehicle-treated S0D1 and vehicle-treated Ntg. Hashtag denotes significance between vehicle-treated S0D1 and PCPZ-/CLP290-treated Ntg. One-way ANOVA with Tukey’s multiple comparison post hoc. Data are presented as mean ± SEM. * P ≤ 0.05, ** P ≤ 0.01, *** P ≤ 0.001, **** P ≤ 0.0001.

### Acute PCPZ administration restores Cl^-^ extrusion capacity in SOD1 and TDP43 mutant mice

In the mature nervous system, KCC2 maintains low intracellular levels of Cl^-^, which in turn influences the strength of GABAergic inhibition ^38,48–50^. To ensure that the effects of the KCC2 enhancers on preservation of motor neurons, motor function and survival was linked to the restoration of KCC2 function and not secondary to other effects of the compounds, we tested the effects of the acute administration of PCPZ on the cellular response in cortical tissue explants from SOD1 mutant mice. While we did not perform motor function assessment in TDP43 mice, we sought to repeat these measures in the TDP43 mutant mouse to confirm that a deficit in KCC2 function and the restoration of said function via administration of PCPZ was not specific only to the SOD1 mutant mouse. To achieve this, we used a Cl^-^ imaging assay designed to measure the rate of transmembrane Cl^-^ transport in response to a step change in extracellular K^+^ concentrations ^28,51^. We found that, in L5-M1 pyramidal neurons in brain slices taken from SOD1 and TDP43 mutant mice, the rate of K^+^-dependent Cl^-^ transport was significantly slower compared to that in slices from non-transgenic littermates (**Fig. 5a & b**). In turn, acute administration of PCPZ significantly enhanced the rate of transport in neurons from both mutant mouse models (**Fig. 5a & b**). These findings indicate that the acute administration of PCPZ was effective at rescuing Cl^-^ extrusion in SOD1 and TDP43 mutant mice.

**FIGURE 5.**
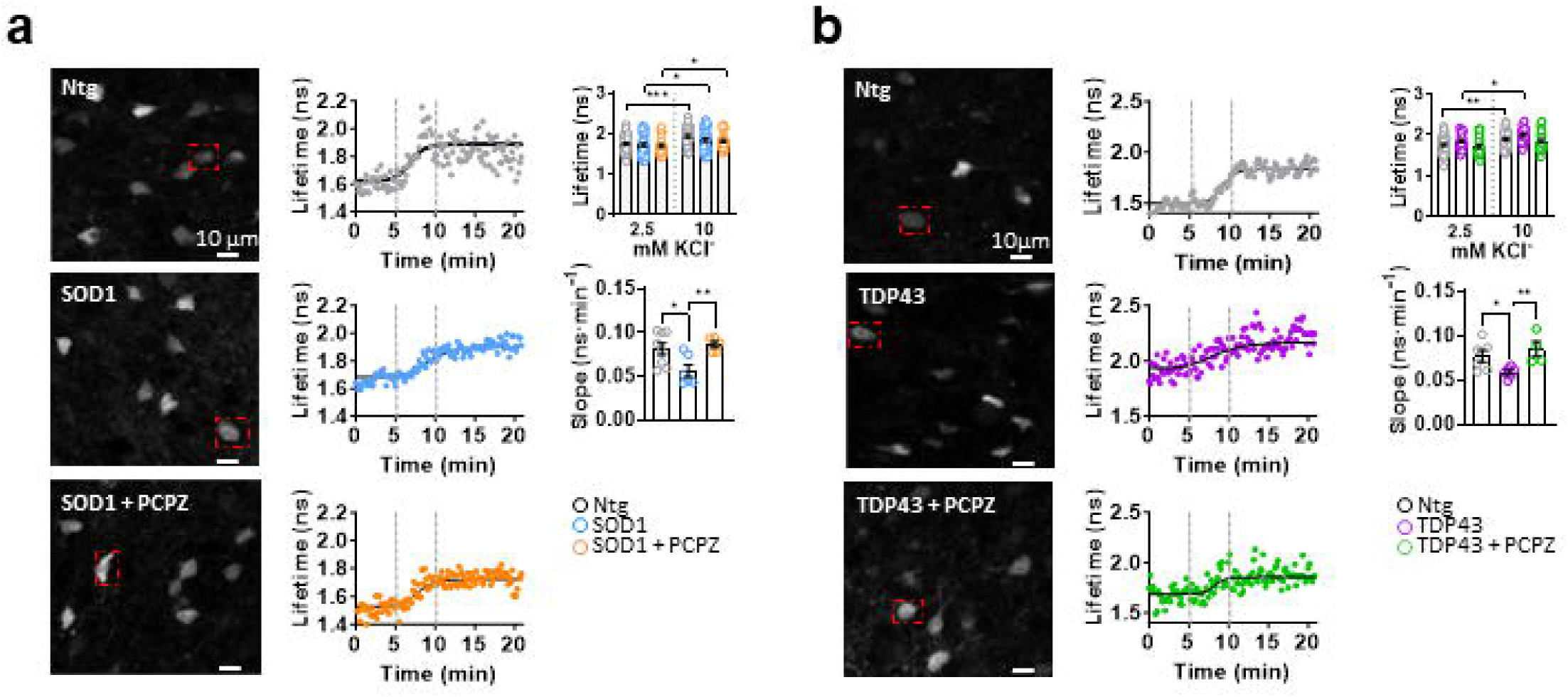
Short-term PCPZ administration restores Cl^-^ extrusion capacity in SOD1 and TDP43 mutant mice. **(a)** (left - right) FLIM images of CaMKIla^+^ neurons expressing SuperClomeleon in the L5-M1 in Ntg, SOD1 and SOD1 slices treated acutely with PCPZ. Example timelapse recording of Ch accumulation in the cell body of selected treated neurons from the ROI shown in images at baseline (2.5 mM KCI) and high KCI (10 mM KCI), onset and timepoints selected to measure slope indicated by the dotted line. Insets show fitted photon distribution histograms at baseline and high KCI. Scale bars: vertical. 50 photons: horizontal. 5 ns. Quantification of the average mean lifetime (paired *t*-test) and mean slope (unpaired *t*-test) measured in all neurons. Ntg, *n = 47 neurons from N = 8 mice;* SOD1, *n = 40 neurons from N = 6 mice:* SOD1+PCPZ, *n = 32 neurons from N* = 5 *mice.* **(b)** Similar to (a), but in TDP43 slices. Ntg, *n = 49 neurons from N = 8 mice.* TDP43. *n = 25 neurons from N = 6 mice;* TDP43+PCPZ. *n = 53 neurons from N = 5 mice.* Data are presented as mean ± SEM. * P ≤ 0.05, ** P ≤ 0 01, *** P ≤ 0.001, **** P ≤ 0.0001.

### Cortical hyperexcitability in awake behaving SOD1 mutant mice is reduced by acute administration of PCPZ

While cortical hyperexcitability has been recognized as a hallmark feature of ALS during the presymptomatic phase of the disease in ALS rodents and patients alike ^1,5,25,52^, there is new evidence suggesting that cortical inhibitory dysfunction directly contributes to this catastrophic phenomenon ^1–3^. To test for a causal link between KCC2 deficits and cortical hyperexcitability, we used a fiber photometry approach to record calcium (Ca^2+^) activity in excitatory neurons within deep layers of the motor cortex in freely moving SOD1 mice (**Fig. 6a)**. Our recordings confirmed, for the first-time, cortical hyperactivity in awake, behaving SOD1 mutant mice as measured by enhanced ongoing fluctuations (mean variance) of the Ca^2+^ signal, compared to non-transgenic littermates ^53^ (**Fig. 6b - d**). Acute treatment of the mice with PCPZ over the course of two days significantly reduced the overt activity in the SOD1 mutant mice, with no significant effect in their control counterparts (**Fig. 6b, c, & e**). These findings establish KCC2 hypofunction as necessary for presymptomatic cortical hyperexcitability.

**FIGURE 6.**
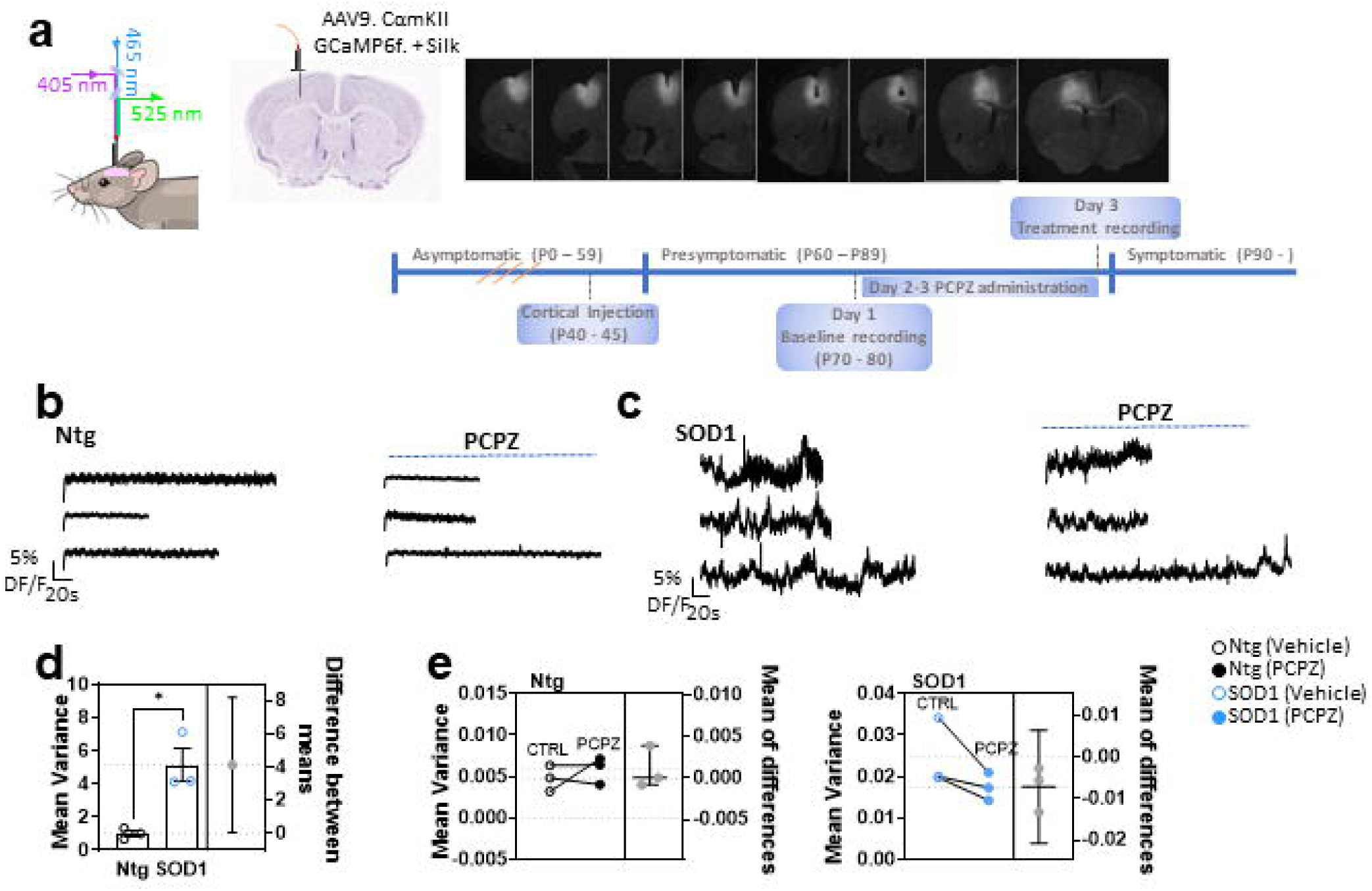
Cortical hyperexcitability in awake behaving SOD1 mutant mice is reduced by short-term administration of PCPZ. **(a)** (left - right) Illustration of light pathway utilized with the Doric photometry device. Location of cannula superimposed onto a coronal slice of the M1. Serial images of the M1 illustrating the high efficiency spread of the AAV9.CamKII GCamp6 Fast construct. Below shows an illustration of experimental design. **(b)** Fiber photometry traces from Ntg mice before (left) and after acute PCPZ treatment (right). **(c)** Similar to (b), but in SOD1 mice. **(d)** Quantification of the trace variance between Ntg and SOD1 mice. Welch’s *t*-test. **(e)** Estimation plots illustrating the effect of PCPZ on cortical excitability in Ntg mice (left) and SOD1 mice (right). *N = 3* for all groups, paired *t*-test .Data are presented as mean ± SEM. *P ≤ 0.05.

### Acute PCPZ administration restores KCC2 expression in homozygous SOD1^G93A^ iPS-derived neurons

To address whether ALS-linked mutations were sufficient to cause a downregulation of KCC2, and to ask weather the administration of PCPZ could be effect in a human cell line, we used human iPSC derived motor neurons with a G93A mutation in the endogenous SOD1 gene. In both two- and four-week-old cultures, we found that KCC2 was decreased in SMI-32^+^-confirmed motor neurons containing the SOD1*G93A relative to isogenic controls (**Fig. 7a –b & Supplemental Fig. 3a - b**). This decrease was specific to KCC2, as we did not observe any difference in MAP2 or β-tubulin expression in either 2- or 4- week cultures (**Fig. 7c – d & Supplemental. Fig. 3c - d**). We next asked whether acute administration of PCPZ was effective at restoring KCC2 membrane expression in these human-derived neurons. To do this, we treated 2- and 4-week-old iPSC cultures with 5 µM PCPZ for either 30-min or 1-h, then quantified KCC2 expression in SMI-32^+^ neurons. We found that a 30-min incubation of the cultures in PCPZ was sufficient to restore KCC2 levels in SMI-32^+^ cells back to that observed in the isogenic controls (**Fig. 7a –b & Supplemental Fig. 3a - b**). This confirms a causal link between the mutation and KCC2 deficits in human motor neurons. In turns, it also validates the potential effectiveness of PCPZ for human therapeutics.

**FIGURE 7.**
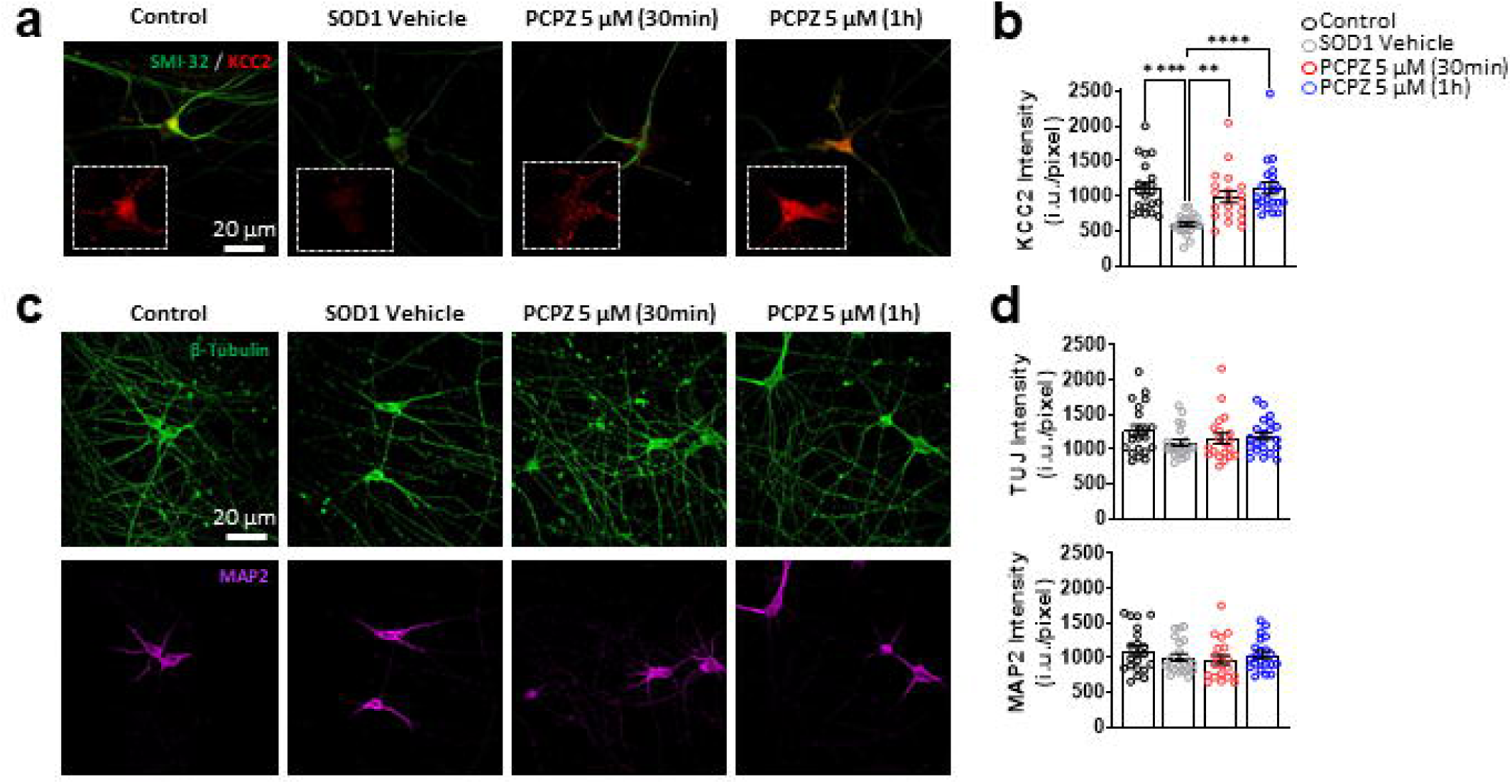
Short-term PCPZ administration restores KCC2 expression at 4 weeks in homozygous SOD1^G93A^ iPS-derived neurons. **(a)** Representative confocal images showing KCC2 labelling in SMI-32^+^ neurons derived from knock-in homozygous SOD1^G93A^ iPSCs at 4 weeks. Cells were treated with either vehicle. 5 μM PCPZ for 30 min or 1h. **(b)** Quantification of mean KCC2 intensity in the cell body across conditions. **(c & d)** Similar to (a & b), but for β-Tubulin (green) and MAP2 (magenta), *n = 18 - 21 cells from 3 independent replicates,* one-way ANOVA with Tukey’s multiple comparison post hoc. Data are presented as mean ± SEM. * P ≤ 0.05, ** P ≤ 0 01, *** P ≤ 0 001, **** P ≤ 0.0001.

## Discussion

Here, we provide evidence that KCC2 membrane expression is decreased and dysfunctional in several key motor areas throughout the corticospinal motor tract in ALS patients and mutant mice during the course of the disease. This in turn, contributes to the onset of motor deficits, altered cortical excitability and ensuing degeneration. We also found increased levels of plasma- and CSF-derived KCC2 can serve as an early biomarker for the disease. Additionally, in line with recent clinical and preclinical reports suggesting that aberrant cortical inhibitory transmission precedes hyperexcitability in both sporadic and familial cases of ALS ^1,3,4^, our results show that the restoration of KCC2 expression and function in presymptomatic ALS mice with either PCPZ or CLP290 is sufficient to decrease cortical hyperexcitability in mice harbouring the SOD1 mutation ^17,18,22,23^.

Persistent deficits in KCC2 expression and function early in development has been associated with changes in neuronal excitation, increased seizure susceptibility, progressive spasticity, low body weight, diminished breathing and learning and memory impairments ^43,54^. Likewise, ALS is associated with motor neuron excitability, motor function deficits and in some cases executive and memory impairments ^55,56^. Thus, an early loss of KCC2 during the juvenile period in SOD1 and TDP43 mutant mice can potentially alter normal brain function by weakening the strength of inhibition, which ultimately leads to changes in CNS network activity. A better understanding of when and where inhibitory deficits occur in the corticospinal motor tract can guide clinicians when assessing patient-specific disease outcome measures and when implementing therapeutics.

Since a definitive diagnostic test for ALS does not exist, patients typically experience delays of ∼ one year between symptom onset and diagnosis ^44–46^. This delay leads to increased psychosocial issues met by both caregivers and patients. Similarly, symptom management, access to therapeutics, and opportunities to enroll in early disease stage clinical trials are also delayed ^45,46^. Therefore, the identification and development of not only a diagnostic, but also prognostic and pharmacodynamic biomarkers, are crucial to increasing the speed at which diagnoses are delivered ^57^. Additionally, while measures of protein concentrations within the CSF tend to have superior sensitivity and specificity, CSF extraction for the purpose of biomarker evaluation is an invasive procedure that may cause hesitation in asymptomatic individuals at risk of developing ALS ^58,59^. Blood-based biomarkers, on the other hand, are often clinically preferential as their collection is both time- and cost-effective and are accompanied by less individual risk. While a larger group of patients across various time points in the disease need to be assessed, our results show promising evidence that changes in KCC2 protein levels can be reliably detected in plasma and CSF extracted from ALS patients, which is paramount to the clinical success of a KCC2-enhacing therapeutic in ALS. While we did not detect a significant change in KCC2 protein levels in the plasma from the SOD1 mouse, it is important to note that these samples were assessed during the presymptomatic phase and thus, represented an early stage in the etiology of the disease when compared to the samples taken from recently diagnosed patients. These results imply that CSF measures may be more sensitive to changes in KCC2 protein levels earlier in the disease progression. It can be speculated that the increase in KCC2 protein levels (or fragments thereof) found in the plasma and CSF of ALS patients mirrors the loss of KCC2 in CNS neurons. While we do not explore the nature of increased KCC2 expression in the plasma and CSF here, it may be the result of protein shedding or degradation from the membrane ^60^. Consistent with this, increase in serum levels of fragmented KCC2 has been observed following CNS trauma ^61^. Regardless of the mechanistic base for enhanced plasma KCC2, our findings establish this as a potential biomarker that is common across multiple genetic forms of the disease.

While it’s clear from the evidence provided here and from others in recent years, that inhibition is altered presymptomatically in ALS, what remains to be demonstrated is how this information can be used to subdue cortical hyperexcitability which has been implicated in the triggering of excitotoxic events that eventually lead to neuronal degeneration ^1–3,6^. Here, we provide evidence, for the first time, that cortical hyperexcitability is present in awake, behaving SOD1 mutant mice and that acute PCPZ administration can effectively counter the exaggerated motor cortical activity. Prochlorperazine is an FDA-approved anti-psychotic and -emetic drug that was identified as a candidate compound to exert effects on KCC2 in a pharmaceutical drug screen ^47^. In line with evidence provided here, others have found that PCPZ, at doses below that of producing anti-psychotic and anti-emetic effects, restores KCC2 expression in spinal cord injury models, and alleviates associated spasticity and recovers muscle activity to baseline ^42,47^. CLP290 has also been shown to significantly enhance and restore Cl^-^ transport in CNS neurons within neuropathic pain and spinal cord injury models that have reduced KCC2 function ^8,39,40,62,63^. KCC2’s confinement to CNS neurons also makes it an ideal target to control Cl^-^ levels while limiting side effects in other organs ^39^. Additionally, with over 60 compounds being evaluated in ALS clinical trials, only three have been approved for clinical use: riluzole, edaravone and AMX0035. In SOD1 mutant mice, riluzole was able to extend the life span by approximately two weeks, while edavarone has negligible effects and AMX0035 prolonged survival by approximately one month ^64–66^. Here, we show that the long-term administration of PCPZ and CLP290 translated into a three to four-week delay in motor symptom onset and extended survival up to two months in the mutant SOD1 mouse. Moreover, our results are the first of their kind to demonstrate the stabilization of weight long-term in SOD1 mutant mice as a result of therapeutic intervention. Weight loss is not only a clinical hallmark feature of the disease, but also indicative of the speed of disease progression and well documented in SOD1 mutant mouse ^67^. Taken together, these findings support that the reversal of KCC2 deficits has the potential to extend lifespan, improve quality of life and lessen caregiver burden. With minimal side effects and being well tolerated for long-term use in humans, PCPZ potentially offers a novel therapeutic avenue for ALS patients.

## Methods

### Animals

Mice were obtained from Jackson Laboratories: SOD1*G93A [B6SJL Tg(SOD1*G93A)1Gur/J] and B6.Cg-Tg(Prnp-TARDBP*A315T)95Balo/J. Wild-type mice were litter- and age-matched. For all behavioural experiments, we used equal proportions of female and male mice with a minimum of 24 mice (12 male: 12 female mice) per condition. Mice were randomly assigned to experimental groups. Mice were housed under a 12/12-h light/dark cycle with ad libitum access to food and water and were randomly allocated to different test groups. All experiments were approved by the committee for animal protection at Université Laval (CPAUL) and are in accordance with the guidelines of the Canadian Council on Animal Care. In our colonies, males from the SOD1*G93A colony reached their natural end points at ∼130 days, while females reached their end points at ∼140 days. These mice typically presented with the onset of significant motor deficits at ∼110. All data are reported in weeks.

### Fluorescent immunolabelling

Mice were transcardially perfused with ice-cold phosphate-buffered saline (PBS), followed by 4% paraformaldehyde (PFA). Brains were extracted and post-fixed in 4% PFA for 16 h at 4°C and cryoprotected in 30% sucrose and stored at −80°C. Slices containing the M1 and/or the ventral horn in the L5 segment of the spinal cord were cut coronally at 30-µm thickness on a Leica cryostat. Free-floating sections containing the M1 and or the L5-SC were rinsed once in PBS for 5 min, followed by two more washes in PBS with 0.1% Triton™ X-100. Slices were then blocked in PBS containing 10% goat serum and 0.1% Triton™ X-100 for 1 h at room temperature, followed by a 16-h incubation in PBS with 0.1% Triton™ X-100 with the following antibodies: mouse monoclonal anti-NeuN (1:500; EMD Millipore, MAB377), rabbit polyclonal anti-KCC2 (1:1000; EMD Millipore; 07-432), rabbit polyclonal anti-MAP2 (1:500; Sigma-Alrich, AB5622), and rabbit polyclonal anti-CTIP2 (1:100, Abcam, ab28448) at 4°C. Finally, slices were incubated in either Alexa Fluor® 488-conjugated goat anti-rabbit antibody (1:500, Thermo Fisher, A27034) for CTIP2, Alexa Fluor® 555-conjugated goat anti-rabbit antibody (1:500, Thermo Fisher, A27039) for KCC2 and MAP2, or Alexa Fluor® 488-conguated goat anti-mouse antibody (1:500, Thermo Fisher, A28175) for NeuN for 2 h at room temperature.

For human tissues, informed consent was obtained from all participants with approval from the Sunnybrook Research Institute Research Ethics Board, in accordance with Clinical Trials Ontario guidelines. Patients were diagnosed at the ALS Clinic at Sunnybrook Health Sciences Centre in Toronto, using the revised El Escorial Criteria. Four SOD1 mutation carriers, 2 C9orf72 and 3 sporadic ALS patients were identified. Human brain slices were paraffin embedded and obtained from Sunnybrook Hospital; all control and ALS cases were collected at the same hospital and treated in the same manner. Briefly, slides were subjected to a series of xylene and ethanol washes to de-paraffin and re-hydrate the tissue, then heated in a steamer for 10 min to ensure antigen-retrieval. Tissue was blocked in TBS containing 5% goat serum for 1.5 h at room temperature, followed by a 38-h incubation in TBS with 0.05% Tween with rabbit polyclonal anti-KCC2 (1:500; EMD Millipore; 07-432) at 4°C. Tissue was then incubated in Alexa Fluor® 555-conjugated goat anti-rabbit antibody (1:500, Thermo Fisher, A27039 for 2 h at room temperature. Finally, slides were quickly incubated in TrueBlack® (1X; biotium; 23007) for 30 s and rinsed 3 times with TBS to limit autofluorescence and lipofuscin.

The use of the IPSCs and IPSC-derived lines was approved by the McGill University Health Centre Research Ethics Board (DURCAN_IPSC/2019-5374). For iPSC cultures, the SOD1 G93A mutation was introduced into a healthy control AIW002-02 iPSC line through CRISPR genome editing, as previously described by ^68^. Cells were plated onto coverslips and treated with either 5 µM PCPZ for 30 min, 1h or vehicle at room temperature. Coverslips were then rinsed once in PBS for 5 min and then blocked in PBS containing 10% goat serum for 20 min at room temperature. Coverslips were then fixed in 4% PFA for 10 min and washed once with PBS for 5 min before being incubated in PBS containing 10% goat serum, 0.1% Triton™ X-100 and the following antibodies; mouse monoclonal anti-SMI-32 (1:100; BioLegend, 801701), rabbit polyclonal anti-KCC2 (1:100; EMD Millipore, 07-432), mouse monoclonal anti-βIII-Tubulin (1:500; EMD Millipore, MAB5564) or chicken polyclonal anti-MAP2 (1:500; EnCor, CPCA-MAP2) for 45 min at room temperature. Finally, coverslips were then washed with PSB for 5 min and then incubated in either, Donkey anti-mouse IgG Dylight 488 (1:500, Abcam, ab96875) for SMI-32 and βIII-Tubulin, Donkey anti-rabbit IgG Dylight 550 (1:500; Abcam, ab96892) for KCC2 or Donkey anti-chicken IgG AlexaFluor 647 (1:500; Jackson Immunoresearch, 703-605-155) for MAP2 for 40 min at room temperature.

### Microscopy and image acquisition

Mounted slices were imaged on a Zeiss LSM710 laser scanning microscope using an MBS 6 405/488/555/639 beam splitter or a Zeiss LSM880 laser scanning microscope using an MBS 7 488/555/639 beam splitter. Images were obtained using a 20×, 40× and 63× oil-immersion objective. Optimal values were selected to limit pixel saturation and avoid photobleaching. The laser power, polarization voltage, filters, dichroic mirrors, and scan speed were kept constant for each unique dataset. In all cases, the L5-M1, L5-VH and Nucleus Accumbens was identified using DAPI staining with the 10× objective and individual cells from this area were chosen at random (experimenter was blinded) with either the 40× or 63× objective. NeuN^+^ and CTIP2^+^ cell bodies were identified, measured, and counted manually in ImageJ with the Fiji plugin. The fluorescence intensity for KCC2 labelling was measured with a custom MATLAB algorithm, referred to as MASC-⫪ which measured the membrane analysis of subcellular intensity, as previously described by ^7,28,69^. Briefly, for each confocal image the membrane of randomly and blindly selected neurons (for which a continuous membrane across the entire cell body circumference was identified) was manually delineated. For each pixel in the ROIs defining individual neurons, the distance to the closest membrane segment was calculated to generate a distance map with the neuronal membrane defined as zero. The mean intensity per pixel and the standard deviation of the KCC2 fluorescence was then computed as a function of the distance to the membrane. This approach provides an unbiased estimate of the membrane expression and excludes any potential effect of neuronal loss in the protein level measurements within the brain regions of interest. MAP2 and β-Tubulin fluorescence was analysed in a similar manner, but fluorescence was measured across the entire cell body. Imaging experiments were performed and analysed in a blinded manner.

### CSF and plasma collection

For presymptomatic collection of plasma and CSF, mice were placed into a stereotaxic apparatus (Stoelting Co.) and maintained on a heating pad under 4% isoflurane anaesthesia. Using a pulled glass capillary, CSF was drawn out with negative pressure from the magna cisterna and stored at -20°C until further use. Following this, mice were decapitated, and blood was immediately collected into K_3_ ETDA coated collection tubes (Thermo Fisher Scientific, 22-030-403) and stored on ice. Collection tubes were then centrifuged at 4°C for 10 minutes at 1,000– 2,000 x g and stored at -20°C until further use.

Plasma and CSF samples from patients were distributed through the NEALS Biofluid Repository (project ID# 2022.02). Informed consent was obtained from all participants with approval from Massachusetts General Hospital Ethics Committee, in accordance guidelines approved in Cross-Sectional ALS Biofluid Biomarker (CABB) Study. Patients were diagnosed by a neurologist using the revised El Escorial Criteria. Four sporadic SOD1-linked ALS patients were identified. Human and mouse Plasma and CSF samples were probed for KCC2 in a mouse SLC12A5 Sandwich ELISA kit (Cedarlane, LS-F65788).

### Drugs

CLP290 was prepared fresh daily by dissolving it in 20% 2-hydroxypropyl-ß-cyclodextrin (HPCD) at 10 mg mL^-^1. Solution was administered at a concentration of 100 mg kg^-1^ via *per os* (*p.o.*) by gavage at the same time daily beginning at the presymptomatic stage (∼P60) until the animal’s endpoint. A 20% HPCD solution was used as the vehicle treatment.

Prochlorperazine (Fisher, P23685G) was dissolved in a solution of 10% DMSO and Lactated Ringer’s saline. For behavioural experiments and survival data, PCPZ was administered at a concentration of 1 mg kg^-1^ daily via *intraperitoneal* (*i.p*.) injection beginning at the presymptomatic stage (∼P60) until the animal’s endpoint. Lactated Ringer’s solution containing 10% DMSO was used as the vehicle treatment. For calcium recordings, PCPZ (1 mg kg^-1^) was administered for 2 days via *i.p.* injection following baseline recording (P70-80).

### Assessment of motor function

Motor performance tests were performed between 90 and 125 days. Natural end point was determined by the animal’s inability to self-correct within 10 s of being placed on its back and a decline in weight of >20% in 72 h. For all experiments, weight measurements were taken twice weekly between 90 to 100 days to establish a baseline, then taken three times per week to evaluate loss of body weight and muscle mass pertaining to disease progression.

To quantitatively assess grip strength, the BIOSEB Grip Strength Test (Model: BIO-GS3) was used, which provided a readout of the maximal peak force of each mouse. Grip strength performance was measured by placing the mouse’s paws on a wire grid until it naturally gripped the bars followed by a gentle pull backwards on the tail. The maximum strength of the grip prior to grip release was recorded. The test was performed in two sessions per week with each session consisting of three trials. The average performance for each session was presented as the average of the three trials/weight.

Rotor-Rod performance was used to assess sensorimotor coordination and motor learning. Mice were placed on a rotating rod that accelerated quickly from 0 to 5 rpm and then gradually from 5 to 20 rpm. Mice were trained for 1 week prior to testing on a rod constantly rotating at 10 rpm. The latency to fall was recorded across two sessions per week, two trials per session; the average of the two trials was presented daily.

### Coherent anti-Stokes Raman spectroscopy

Spinal cords were extracted and post-fixed in 4% PFA/glutaraldehyde solution for 72 h from the same mice that were transcardially perfused for fluorescent immunolabelling. Cords were removed from spinal columns and L5 roots were dissected away in PBS. Ventral roots from the L5 spinal segment were embedded in 4% agarose (made in PBS). Coronal slices (350 μm) were prepared using a Leica vibratome and mounted just prior to imaging. The Coherent anti-Stokes Raman spectroscopy (CARs) setup used was based on a custom-build, video-rate, laser scanning microscope and two laser sources. An 80 MHz, 7 ps mode-locked laser (Nd: YVO4, High-Q Laser) delivered a Stokes pulse at 1064 nm, and its second harmonic synchronously pumps an optical parametric oscillator (OPO) (Levante Emerald, APE), generating a pump beam which was set at 817 nm to probe CH2 stretching bands in myelin (2845/cm). The pump and the Stokes beams were recombined spatially using a dichroic mirror (FF875 Di01) and temporally using a delay line. The laser beams were focused on the sample using a high-numerical aperture water-immersion objective (60×, 1.2NA, UIS-UPLAPO, Olympus). Images were acquired by raster beam scanning of the sample (gold-coated polygonal mirror for the fast axis, Lincoln Laser, DT-36-290-025, a galvanometer mirror for the slow axis, Cambridge Technology 6240H) at a rate of 30 images/s (2252 × 500 pixels, scan range 300 × 150 µm) and with an average power at the focal spot of 10 mW. The backscattered anti-Stokes signal (662 nm) is separated from the excitation beam by a dichroic long-pass filter (Semrock, FF735-Di01) and a short-pass filter (Semrock, FF01-750/SP-25) and collected in the epi-direction by a red-sensitive photomultiplier tube (Hamamatsu, R3896). Motor axons were identified and counted manually using ImageJ with the Fiji plugin.

### Surgical procedures

Stereotaxic surgery was conducted on mice placed into a stereotaxic apparatus (Stoelting Co.) and maintained on a heating pad under 1.5 - 4% isoflurane anaesthesia. For *ex vivo* Cl^-^ imaging experiments, AAV2/9.CaMKIIa.Superclomeleon (purified and produced by the Canadian Neurophotonics Platform Viral Vector Core; RRID:SCR_016477) was bilaterally infused into the M1 (rostro-caudal: +1.5, dorso-ventral: −1.6, latero-medial: ±1.5 from Bregma) of SOD1, T43 and their non-transgenic littermates at a volume of 100 nL. All infusions were made using a glass capillary and a NANOLITER2020 injector (World Precision Instruments LLC). Mice were returned to their cages and housed with their littermates. Imaging started 4 weeks after viral injection to ensure viral expression.

For *in vivo* Ca^2+^ photometry experiments, 1.6 mm mono fiberoptic cannulas (MFC_400/430-0.66_1.6mm_MF1.25_FLT Mono Fiberoptic Cannula, Doric) were coated with 200 nL of a 1:1 mixture of 5% silk fibers and AAV9.CaMKII.GCaMP6f.WPRE.SV40 (Addgene viral prep # 100834-AAV9), as previously described by ^70^. Cannulas were then implanted unilaterally into the M1 (rostro-caudal: +1.5, dorso-ventral: −1.6, latero-medial: ±1.5 from Bregma) between P40 and 45 and secured to the skull using C&B Metabond and an anchoring screw contralateral to the cannula. Mice were single-housed with all supplemental enrichment material removed. Experiments began a minimum of 4 weeks post-implantation to allow for viral expression.

### Brain slice preparation and FLIM

Brains were rapidly removed after decapitation and placed into ice-cold cutting saline containing the following (in mM): 205 sucrose, 2.5 KCl, 1 CaCl_2_, 2 MgCl_2_, 1.25 NaH_2_PO_4_, 25 NaHCO_3_, 25 Glucose, 0.4 Na-Ascorbate and 3 Na-Pyruvate in double-distilled water (ddH2O), pH 7.4 with NaOH, osmolarity 300–305 mOsm saturated with 95% O_2_/5% CO_2_. Coronal brain slices (300 μm) were prepared from SOD1, TDP43 and non-transgenic littermates using a Leica vibratome. Slices were then incubated for 30 min at 32 °C in a 1:1 mix of cutting saline and aCSF containing (in mM); 126 NaCl, 5 KCl, 2 CaCl_2_, 2 MgCl_2_, 10 Glucose, 26 NaHCO_3_ and 1.25 NaH_2_PO_4_; pH 7.35 and 295 mOsm, followed by 30 min recovery at room temperature in oxygenated aCSF. All chemicals were purchased from Sigma, unless otherwise specified.

For all FLIM experiments, coronal brain slices expressing SuperClomeleon ^71^ were prepared from presymptomatic mice (P65-P75) and transferred into the perfusion chamber and perfused with aCSF at a rate of 2 mL min^-1^ at room temperature. All recordings were obtained from the L5-M1. The slices were incubated 10 minutes in aCSF containing low extracellular KCl (2.5mM) and 5 μM bicuculline, 10 μM CQNX and 50 μM APV. Fluorescence lifetime values of CaMKIIa^+^ neurons were acquired in aCSF containing low extracellular KCl (2.5 mM) to measure the baseline, followed by aCSF containing low extracellular KCl (2.5 mM) and 5 μM bicuculline, 10 μM CQNX and 50 μM APV; and aCSF containing high extracellular KCl (10 mM) with bicuculline, CQNX and APV. For PCPZ experiments, fluorescence lifetime values were acquired under the same conditions following a 1 h incubation in 10 µM PCPZ or vehicle diluted in aCSF for 1 h at room. Each FLIM image corresponds to 10 seconds acquisitions. Baseline recordings were acquired for 2.5 mins in the presence of aCSF containing low extracellular KCl (2.5 mM); then 6 mins recordings were acquired in the presence aCSF with low extracellular KCl (2.5 mM) plus blockers, followed by a 15 min recording in the presence of aCSF with high extracellular KCl (10 mM) plus blockers.

Fluorescence lifetime imaging microscopy was performed as previously described by ^11^ using a Zeiss 880 laser-scanning microscope coupled with a TCSPC module PMC-100-1 DETECTOR (Becker & Hickl GmbH). A 80 MHz femtosecond-pulsed Ti-Sapphire laser set at 800 nm and used for excitation. FLIM images (1 per 10s) were acquired with a 20x Plan-Apochromat water immersion objective (Zeiss, 1.0 NA), and FLIM images were generated using a TCSPC module PMC-100-1 detector (Becker & Hickl GmbH) with a bandpass filter (Chroma, ET473/24m) to collect only the donor emissions.

For each acquisition, neuronal cell bodies were delineated into individual regions of interest (ROIs). The instrument response function (IRF) was acquired by using an 80 nm gold nanoparticle to generate a second-harmonic signal. Using a custom script, the FLIM histogram was generated by pooling all the pixels in the ROIs for each timepoint. Every time point (10 seconds) of the experiment was fitted using a mono-exponential decay, and the fluorescence lifetime (*τ*) was estimated. For each neuron analyzed, a Hill-Slope was fitted into the temporal evolution of lifetimes to estimate the initial and final lifetime values (for 2.5 mM and 10 mM KCl aCSF, respectively). The maximum slope of the lifetime (units of ns per min) was estimated upon 10 mM KCl application using a plateau followed by an exponential plateau function. Neurons that did not have enough photons in the FLIM acquisition (300 photons), a stable lifetime during the 5 minutes before the potassium increase, or did not respond, meaning no significant increase of lifetime was observed after the potassium increase, were excluded from the analysis.

### Calcium Photometry

SOD1*G93A mice and their non-transgenic littermates implanted with mono fiber-optic cannulas and expressing AAV9.CaMKII.GCaMP6f. were fixed to a 400 µm core diameter low autofluorescence mono fiber-optic patch cord with a rotary joint which was connected to a integrated fluorescence mini cube (excitation wavelengths 405 and 465 nm) and placed into a clean empty cage. Mice were allowed 5 min to acclimatize to the cage before beginning acquisition. Five-minute recordings were acquired from mice (P70 -80) which were allowed to freely move about in an empty cage with a food grid above which served as a baseline. Mice were then returned to their home cages and received daily i.p. injections of PCPZ (1 mg kg^-1^) for a period of 2 days after which they were returned to a clean empty cage and 5 min recordings were acquired once again. Two traces were analyzed used a custom MATLAB algorithm. The first one corresponds to the trace recorded using the 465 nm excitation while the second trace was recorded using the 405 nm which corresponds to the isosbestic point of GCaMP that provides a signal that is calcium independent. We first found the linear transformation that maximize the overlap between the reference to the signal trace. We subtracted the trace recorded using the 465 nm excitation from the adjusted reference trace recorded using the 405 nm. Finally, we divided the resulting trace by the adjusted reference trace to obtain the delta F over F trace (DF/F) from which the mean variance was calculated. Each mouse served as its own control.

### Statistics

All data are presented as mean ± standard error of the mean (SEM), and each datum point represents an individual animal or experiment. Tests of statistical difference were performed with GraphPad Prism 10 software using unpaired and paired t-tests (two-sided) or one-way ANOVA with Sidak’s multiple comparisons or Bonferroni post hoc test. Time course behavioural experiments were analysed using a two-way ANOVA with Sidak’s multiple comparisons test. Sample sizes are consistent with those reported in similar studies and provide sufficient power to detect changes with the appropriate statistical analysis. Data used for statistical comparison fit a Gaussian (normal) distribution, and variances were equivalent between groups. In cases where the normality did not hold, a non-parametric test (Mann-Whitney) was used for independent samples. See manuscript for specifics. For all experiments, a criterion α-level was set at 0.05 (not significant P > 0.05, *P ≤ 0.05, **P ≤ 0.01 ***P ≤ 0.001 ****P ≤ 0.0001). For all behavioural and Ca^2+^ photometry experiments each n-value represents an individual animal. For *ex vivo* Cl^-^ experiments, n-values represent individual recordings, which in some instances were made from the same animal; when n-values represent individual recordings, the number of animals those recordings were made from are clearly indicated in the associated figure legend. For immunofluorescence experiments, cell counts were made from multiple slices from the same animal, and these were averaged and reported as one n-value.

## References

1. Khademullah, C. S. et al. Cortical interneuron-mediated inhibition delays the onset of amyotrophic lateral sclerosis. Brain 143, 800–810 (2020).

2. Van Den Bos, M. A. J., et al. Imbalance of cortical facilitatory and inhibitory circuits underlies hyperexcitability in ALS. Neurology 91, E1669–E1676 (2018).

3. Zhang, W. et al. Hyperactive Somatostatin Interneurons Contribute to Excitotoxicity in Neurodegenerative Disorders. Nat Neurosci 19, 557 (2016).

4. Geevasinga, N. et al. Cortical Function in Asymptomatic Carriers and Patients With C9orf72 Amyotrophic Lateral Sclerosis. JAMA Neurol 72, 1268–74 (2015).

5. Wainger, B. J. et al. Intrinsic membrane hyperexcitability of amyotrophic lateral sclerosis patient-derived motor neurons. Cell Rep 7, 1–11 (2014).

6. Vucic, S. & Kiernan, M. C. Novel threshold tracking techniques suggest that cortical hyperexcitability is an early feature of motor neuron disease. Brain 129, 2436–2446 (2006).

7. Lorenzo, L. E. et al. Enhancing neuronal chloride extrusion rescues α2/α3 GABAA-mediated analgesia in neuropathic pain. Nature Communications 2020 11:1 11, 1–23 (2020).

8. Chen, B. et al. Reactivation of Dormant Relay Pathways in Injured Spinal Cord by KCC2 Manipulations. Cell 174, 521 (2018).

9. Dargaei, Z. et al. Restoring GABAergic inhibition rescues memory deficits in a Huntington’s disease mouse model. Proc Natl Acad Sci U S A 115, E1618–E1626 (2018).

10. Merner, N. D. et al. Regulatory domain or CpG site variation in SLC12A5, encoding the chloride transporter KCC2, in human autism and schizophrenia. Front Cell Neurosci 9, (2015).

11. Keramidis, I. et al. Restoring neuronal chloride extrusion reverses cognitive decline linked to Alzheimer’s disease mutations. Brain (2023) doi:10.1093/BRAIN/AWAD250.

12. Masrori, P. & Van Damme, P. Amyotrophic lateral sclerosis: a clinical review. Eur J Neurol 27, 1918–1929 (2020).

13. Cleveland, D. W. & Rothstein, J. D. From Charcot to Lou Gehrig: deciphering selective motor neuron death in ALS. Nat Rev Neurosci 2, 806–819 (2001).

14. Román, G. C. Neuroepidemiology of amyotrophic lateral sclerosis: clues to aetiology and pathogenesis. J Neurol Neurosurg Psychiatry 61, 131–137 (1996).

15. Chiò, A. et al. Global epidemiology of amyotrophic lateral sclerosis: a systematic review of the published literature. Neuroepidemiology 41, 118–30 (2013).

16. Wijesekera, L. C. & Leigh, P. N. Amyotrophic lateral sclerosis. Orphanet J Rare Dis 4, 3 (2009).

17. Zarei, S. et al. A comprehensive review of amyotrophic lateral sclerosis. Surg Neurol Int 6, 171 (2015).

18. Renton, A. E., Chiò, A. & Traynor, B. J. State of play in amyotrophic lateral sclerosis genetics. Nat Neurosci 17, 17–23 (2014).

19. Fogarty, M. J. et al. Cortical synaptic and dendritic spine abnormalities in a presymptomatic TDP-43 model of amyotrophic lateral sclerosis. Sci Rep 6, (2016).

20. Reale, L. A. et al. Pathologically mislocalised TDP-43 in upper motor neurons causes a die-forward spread of ALS-like pathogenic changes throughout the mouse corticomotor system. Prog Neurobiol 102449 (2023) doi:10.1016/J.PNEUROBIO.2023.102449.

21. Brunet, A., Stuart-Lopez, G., Burg, T., Scekic-Zahirovic, J. & Rouaux, C. Cortical Circuit Dysfunction as a Potential Driver of Amyotrophic Lateral Sclerosis. Front Neurosci 14, (2020).

22. Menon, P., Kiernan, M. C. & Vucic, S. Cortical hyperexcitability precedes lower motor neuron dysfunction in ALS. Clinical Neurophysiology 126, 803–809 (2015).

23. Vucic, S., Nicholson, G. A. & Kiernan, M. C. Cortical hyperexcitability may precede the onset of familial amyotrophic lateral sclerosis. Brain 131, 1540–1550 (2008).

24. Menon, P. et al. Cortical hyperexcitability evolves with disease progression in ALS. Ann Clin Transl Neurol 7, (2020).

25. van Zundert, B. et al. Neonatal Neuronal Circuitry Shows Hyperexcitable Disturbance in a Mouse Model of the Adult-Onset Neurodegenerative Disease Amyotrophic Lateral Sclerosis. Journal of Neuroscience 28, 10864–10874 (2008).

26. Fuchs, A. et al. Downregulation of the potassium chloride cotransporter KCC2 in vulnerable motoneurons in the SOD1-G93A mouse model of amyotrophic lateral sclerosis. J Neuropathol Exp Neurol 69, 1057–70 (2010).

27. Blaesse, P., Airaksinen, M. S., Rivera, C. & Kaila, K. Cation-chloride cotransporters and neuronal function. Neuron 61, 820–38 (2009).

28. Ferrini, F. et al. Differential chloride homeostasis in the spinal dorsal horn locally shapes synaptic metaplasticity and modality-specific sensitization. Nat Commun 11, (2020).

29. Gruzman, A. et al. Common molecular signature in SOD1 for both sporadic and familial amyotrophic lateral sclerosis. Proc Natl Acad Sci U S A 104, 12524–12529 (2007).

30. Julien, J.-P. & Kriz, J. Transgenic mouse models of amyotrophic lateral sclerosis. Biochimica et Biophysica Acta (BBA) - Molecular Basis of Disease 1762, 1013–1024 (2006).

31. Jonsson, P. A. et al. Disulphide-reduced superoxide dismutase-1 in CNS of transgenic amyotrophic lateral sclerosis models. doi:10.1093/brain/awh704.

32. Grad, L. I., Rouleau, G. A., Ravits, J. & Cashman, N. R. Clinical Spectrum of Amyotrophic Lateral Sclerosis (ALS). Cold Spring Harb Perspect Med 7, (2017).

33. Philips, T. & Rothstein, J. D. Rodent Models of Amyotrophic Lateral Sclerosis. Current protocols in pharmacology */ editorial board,* S.J. Enna *(editor-in-chief) … [*et al.] 69, 5.67.1 (2015).

34. Ling, S. C., Polymenidou, M. & Cleveland, D. W. Converging mechanisms in ALS and FTD: disrupted RNA and protein homeostasis. Neuron 79, 416–438 (2013).

35. Neumann, M. et al. Ubiquitinated TDP-43 in frontotemporal lobar degeneration and amyotrophic lateral sclerosis. Science 314, 130–133 (2006).

36. Chen, S., Sayana, P., Zhang, X. & Le, W. Genetics of amyotrophic lateral sclerosis: an update. Mol Neurodegener 8, (2013).

37. Bäumer, D., Talbot, K. & Turner, M. R. Advances in motor neurone disease. J R Soc Med 107, 14– 21 (2014).

38. Payne, J. A., Stevenson, T. J. & Donaldson, L. F. Molecular characterization of a putative K-Cl cotransporter in rat brain: A neuronal-specific isoform. Journal of Biological Chemistry 271, 16245–16252 (1996).

39. Gagnon, Martin, Bergeron, Marc, Lavertu, Guillaume, Castonguay, Annie, Tripathy, S. Chloride extrusion enhancers as novel therapeutics for neurological diseases. Nat Med 19, 1524–1528 (2013).

40. Donneger, F. Characterization and therapeutic interest of potassium-chloride co-transporter type 2 (KCC2) potentiators in temporal lobe epilepsies. (Sorbonne University, 2022).

41. Chen, B. et al. Reactivation of Dormant Relay Pathways in Injured Spinal Cord by KCC2 Manipulations. Cell 174, 521–535.e13 (2018).

42. Bilchak, J. N., Yeakle, K., Caron, G., Malloy, D. & Côté, M. P. Enhancing KCC2 activity decreases hyperreflexia and spasticity after chronic spinal cord injury. Exp Neurol 338, 113605 (2021).

43. Tornberg, J., Voikar, V., Savilahti, H., Rauvala, H. & Airaksinen, M. S. Behavioural phenotypes of hypomorphic KCC2-deficient mice. European Journal of Neuroscience 21, 1327–1337 (2005).

44. Xu, R. S. & Yuan, M. Considerations on the concept, definition, and diagnosis of amyotrophic lateral sclerosis. Neural Regen Res 16, 1723 (2021).

45. Paganoni, S. et al. Diagnostic timelines and delays in diagnosing amyotrophic lateral sclerosis (ALS). Amyotroph Lateral Scler Frontotemporal Degener 15, 453 (2014).

46. Richards, D., Morren, J. A. & Pioro, E. P. Time to diagnosis and factors affecting diagnostic delay in amyotrophic lateral sclerosis. J Neurol Sci 417, (2020).

47. Liabeuf, S. et al. Prochlorperazine Increases KCC2 Function and Reduces Spasticity after Spinal Cord Injury. J Neurotrauma 34, 3397–3406 (2017).

48. Sigel, E. & Steinmann, M. E. Structure, function, and modulation of GABA(A) receptors. J Biol Chem 287, 40224–31 (2012).

49. Ben-Ari, Y., Cherubini, E., Corradetti, R. & Gaiarsa, J. L. Giant synaptic potentials in immature rat CA3 hippocampal neurones. J Physiol 416, 303–25 (1989).

50. Kaila, K. & Voipio, J. Postsynaptic fall in intracellular pH induced by GABA-activated bicarbonate conductance. Nature 330, 163–165 (1987).

51. Doyon, N. et al. Efficacy of Synaptic Inhibition Depends on Multiple, Dynamically Interacting Mechanisms Implicated in Chloride Homeostasis. PLoS Comput Biol 7, e1002149 (2011).

52. Kuo, J. J. et al. Hyperexcitability of cultured spinal motoneurons from presymptomatic ALS mice. J Neurophysiol 91, 571–5 (2004).

53. Muir, J. et al. In Vivo Fiber Photometry Reveals Signature of Future Stress Susceptibility in Nucleus Accumbens. Neuropsychopharmacology 2018 43:2 43, 255–263 (2017).

54. Hübner, C. A. et al. Disruption of KCC2 reveals an essential role of K-Cl cotransport already in early synaptic inhibition. Neuron 30, 515–524 (2001).

55. Barulli, M. R. et al. Episodic memory and learning rates in amyotrophic lateral sclerosis without dementia. Cortex 117, 257–265 (2019).

56. Machts, J. et al. Memory deficits in amyotrophic lateral sclerosis are not exclusively caused by executive dysfunction: a comparative neuropsychological study of amnestic mild cognitive impairment. BMC Neurosci 15, 83 (2014).

57. Staats, K. A., Borchelt, D. R., Tansey, M. G. & Wymer, J. Blood-based biomarkers of inflammation in amyotrophic lateral sclerosis. Mol Neurodegener 17, 1–19 (2022).

58. Vernau, W., Vernau, K. A. & Sue Bailey, C. Cerebrospinal Fluid. Clinical Biochemistry of Domestic Animals 769 (2008) doi:10.1016/B978-0-12-370491-7.00026-X.

59. Sturmey, E. & Malaspina, A. Blood biomarkers in ALS: challenges, applications and novel frontiers. Acta Neurol Scand 146, 375 (2022).

60. Lichtenthaler, S. F., Lemberg, M. K. & Fluhrer, R. Proteolytic ectodomain shedding of membrane proteins in mammals—hardware, concepts, and recent developments. EMBO J 37, 99456 (2018).

61. Jantzie, L. L., Winer, J. L., Corbett, C. J. & Robinson, S. Erythropoietin Modulates Cerebral and Serum Degradation Products from Excess Calpain Activation following Prenatal Hypoxia-Ischemia. Dev Neurosci 38, 15 (2016).

62. Ostroumov, A. et al. Stress Increases Ethanol Self-Administration via a Shift toward Excitatory GABA Signaling in the Ventral Tegmental Area. Neuron 92, 493–504 (2016).

63. Sullivan, B. J., Kipnis, P. A., Carter, B. M., Shao, L. R. & Kadam, S. D. Targeting ischemia-induced KCC2 hypofunction rescues refractory neonatal seizures and mitigates epileptogenesis in a mouse model. Sci Signal 14, eabg2648 (2021).

64. Gurney, M. E., Fleck, T. J., Himes, C. S. & Hall, E. D. Riluzole preserves motor function in a transgenic model of familial amyotrophic lateral sclerosis. Neurology 50, 62–6 (1998).

65. Ito, H. et al. Treatment with edaravone, initiated at symptom onset, slows motor decline and decreases SOD1 deposition in ALS mice. Exp Neurol 213, 448–455 (2008).

66. Ryu, H. et al. Sodium phenylbutyrate prolongs survival and regulates expression of anti-apoptotic genes in transgenic amyotrophic lateral sclerosis mice. J Neurochem 93, 1087–1098 (2005).

67. Maksimovic, K., Youssef, M., You, J., Sung, H. K. & Park, J. Evidence of Metabolic Dysfunction in Amyotrophic Lateral Sclerosis (ALS) Patients and Animal Models. Biomolecules 2023, Vol. 13, *Page* 863 **13**, 863 (2023).

68. Deneault, E. et al. A streamlined CRISPR workflow to introduce mutations and generate isogenic iPSCs for modeling amyotrophic lateral sclerosis. Methods 203, 297–310 (2022).

69. Dedek, A. et al. Sexual dimorphism in a neuronal mechanism of spinal hyperexcitability across rodent and human models of pathological pain. Brain 145, 1124–1138 (2022).

70. Jackman, S. L. et al. Silk Fibroin Films Facilitate Single-Step Targeted Expression of Optogenetic Proteins. Cell Rep 22, 3351 (2018).

71. Grimley, J. S. et al. Visualization of Synaptic Inhibition with an Optogenetic Sensor Developed by Cell-Free Protein Engineering Automation. The Journal of Neuroscience 33, 16297 (2013).

