## Supplementary figures and images for "KCC2 as a novel biomarker and therapeutic target for motoneuron degenerative disease"

### Supplemental Figure 1

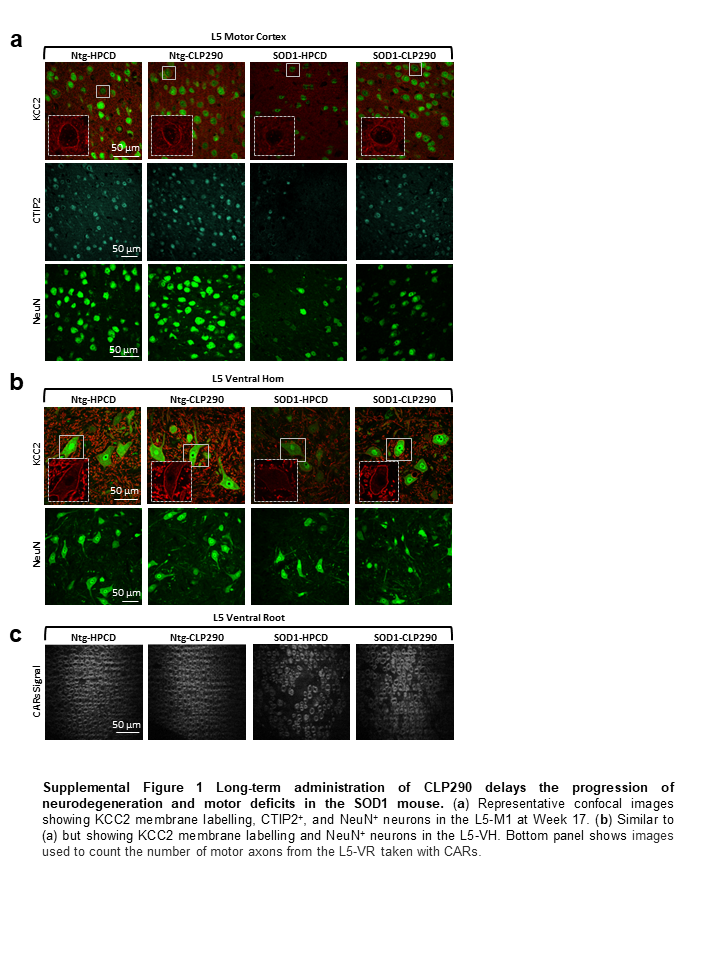

### Supplemental Figure 2

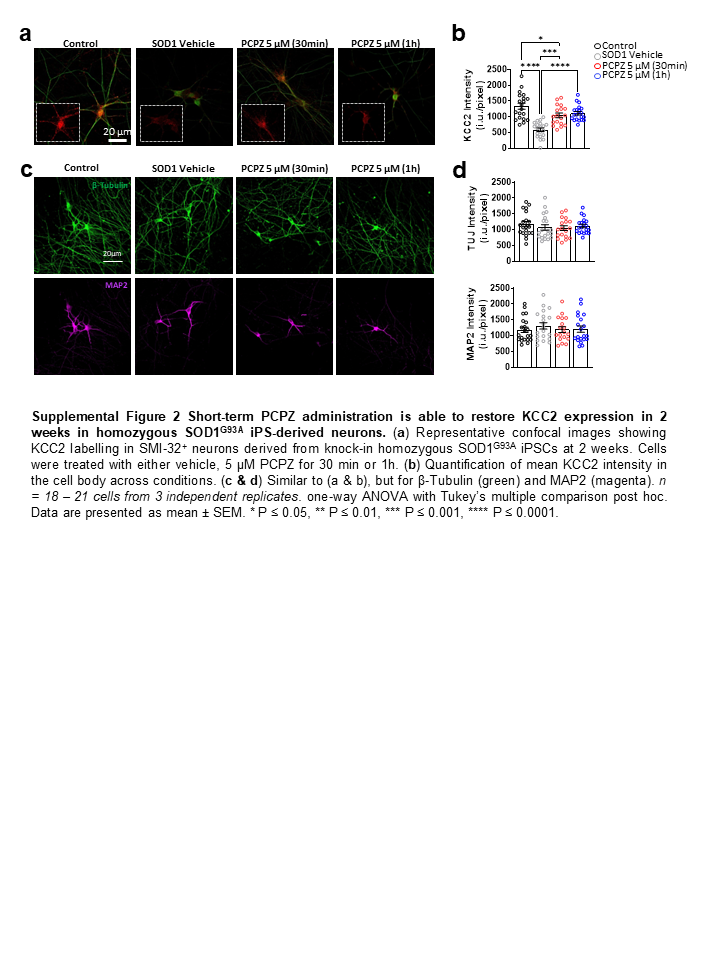
